# 16S-23S-5S rRNA Database: a comprehensive integrated database of archaeal and bacterial rRNA sequences, alignments, intragenomic heterogeneity, and secondary structures

**DOI:** 10.1101/2025.06.23.661116

**Authors:** Jaimin Chodvadiya, Gyanesh Shukla, Gaurav Sharma

## Abstract

The 16S-23S-5S ribosomal RNA (rRNA) gene operon is a cornerstone of ribosomal architecture and function, making it an indispensable molecular marker for microbial taxonomy and phylogenetics. Despite the extensive development of 16S rRNA gene databases, a significant void persists in comprehensive resources that integrate all three rRNA genes (16S, 23S, and 5S), their precise alignments, intragenomic heterogeneity, and intricate structural details. To address this critical research gap, we introduce the 16S-23S-5S rRNA Database, a meticulously curated and intuitive platform encompassing around 5000 complete bacterial and archaeal genomes. This database provides granular operon-level annotations, robust alignment data, quantitative copy number statistics, percent identity matrices, and predicted secondary structures for each rRNA component. Our rigorous methodology integrates covariance models from RFAM with BLAST-based analyses for sequence comparisons, Clustal Omega for multiple sequence alignment, and Circos for genomic visualization, thereby elucidating the genomic architecture, sequence diversity, and structural conservation of rRNA genes. Crucially, the database systematically accounts for intragenomic heterogeneity, a factor paramount for accurate microbial classification and the inference of evolutionary trajectories. Our analysis reveals prevalent phyla such as Pseudomonadota (1,891 genomes), Actinomycetota (846 genomes), and Bacillota (773 genomes), with Bacillota exhibiting the highest average rRNA gene copy numbers (e.g., 16S = 6.64 ± 3.3, 23S = 6.63 ± 3.3, and 5S = 6.8 ± 3.3). Across all bacterial genomes, the 16S, 23S, and 5S rRNA genes exhibited average copy numbers of 4.25 ± 2.88, 4.24 ± 2.89, and 4.44 ± 3.08, respectively. The responsive web interface, engineered with HTML5, JavaScript, and Flask, incorporates user feedback mechanisms to enhance usability. By integrating these comprehensive elements, the 16S-23S-5S rRNA Database not only fills a substantial lacuna in the field of rRNA data analysis but also establishes a foundational resource for large-scale investigations in microbial systematics, genome evolution, and ribosomal biology, benefiting taxonomists, microbial genomics, and microbiome researchers alike.

**Availability and implementation: The** 16S-23S-5S rRNA Database is available as an open-access database at https://project.iith.ac.in/sharmaglab/rrnadatabase/.

**Importance:** Understanding the diversity of microbes is essential for research in health, environment, and biotechnology. Ribosomal RNA (rRNA) genes, especially the 16S gene, are commonly used to classify microbes, but studies often ignore the full set of rRNA genes (16S, 23S, and 5S) together. Our work bridges this gap by creating the first comprehensive database that includes all three rRNA genes from around 5,000 microbial genomes. It offers detailed annotations, sequence alignments, intragenomic heterogeneity, and structural insights, helping scientists more accurately classify microbes and trace their evolution. This resource is designed to be easy to use and will support a wide range of microbiological research.

## Introduction

The 16S ribosomal RNA (rRNA) gene has long served as a foundational marker for microbial taxonomy, systematics, and phylogenetics. Since Carl Woese’s pioneering use of the 16S rRNA sequence to redefine the tree of life into three domains, i.e., Archaebacteria, Eubacteria, and Eukaryotes [1], this gene has been at the heart of molecular microbial ecology. Its conserved sequence and structure interspersed with hypervariable regions allow for both universal detection and taxonomic discrimination of microbial taxa, enabling its widespread adoption across environmental and clinical studies [2]. Advancements in PCR and sequencing technologies, particularly high-throughput sequencing, in the last two decades, have made it possible to investigate microbial communities across diverse ecological and clinical niches and the majority of such studies are using 16S rRNA gene amplicons [12]. This has led to an explosion of sequencing data, enabling the rise of microbiome research and studies in microbial ecology, evolution, human health, agriculture, and more. The centrality of the 16S rRNA gene in microbial research has been solidified by the development of curated reference databases and computational pipelines, further standardizing and streamlining microbial classification efforts globally. The development of large-scale 16S-based resources such as SILVA [3], the Ribosomal Database Project (RDP) [4], Greengenes [5], MIrROR [6], GROND [7], and more recently Greengenes2 [8], has greatly facilitated high-throughput microbial community profiling and classification. Since their inception, these databases have also been well integrated into numerous pipelines, tools, and taxonomic frameworks as curated reference sequences essential for the homology-based classification of microbiomes.

Despite its immense utility in the taxonomic classification, this ∼1500 bp long 16S rRNA gene is only one part of the ribosomal RNA operon. In prokaryotes, the ribosome comprises three rRNA components, i.e., 16S (∼1500 bp long and a part of small ribosomal subunit), 23S and 5S (∼2900 bp and ∼120 bp long, respectively, and part of large ribosomal subunit) rRNAs, encoded within a single operon that is often co-transcribed, majorly alongside tRNA genes [9]. These operons are processed by a complex set of nucleases and RNA-modifying enzymes to yield the mature rRNAs required for ribosome assembly and function [10]. During active exponential growth phases, transcription of rRNA genes accounts for over half of the cell’s total transcriptional output, underlining their central role in cellular physiology [11]. The evolutionary constraints on rRNAs have rendered them highly conserved molecules yet interspersed with regions of variability, especially within the 16S rRNA, that can inform phylogenetic and functional distinctions.

Due to the ease of sequencing data available, 16S rRNA-based community analysis became an important tool for studying microbes in different environments [12]. However, a key complication arises from gene copy number (GCN) variability, as many bacterial and archaeal genomes harbor multiple rRNA operons with sequence variants that can exhibit intragenomic heterogeneity [13, 14]. This heterogeneity can lead to the misclassification of a single organism’s rRNA copies into different taxonomic clusters, thereby creating artificial inflation of microbial diversity in amplicon-based studies. A previous study on complete genomes of bacteria and archaea reported intragenomic heterogeneity in 952 genomes, of which 12.5% divergence reported being above the 1% level [15]. A recent study analyzing 24,248 complete genomes, reported heterogeneity as high as 27.9 % in *Listeria monocytogenes* 10-092876-1155 LM6, overall suggesting that the presence of such multiple 16S rRNA variants can result in their classification into distinct sequence clusters, thus potentially overestimating microbial diversity [14]. Therefore, systematic accounting intra-genomic sequence diversity in reference databases is much needed.

In the past three decades, several 16S gene and operon-oriented databases have been developed, which have proved instrumental in curating rRNA sequences and have been immensely supporting taxonomic classification. Among the most widely used resources, public databases such as SILVA [3], the Ribosomal Database Project (RDP) [4], Greengenes [5], MIrROR [6], GROND [7], and more recently Greengenes2[8], provide high-quality rRNA sequences with curated taxonomy and additionally serves as a reference for the classification of 16S rRNA sequences **(Table 1)**. Among these, SILVA and RDP databases contain millions of high-quality rRNA sequence data with curated taxonomy that serves as a reference for taxonomic classification of high throughput sequencing data [3, 4]. Despite their contributions, these resources are predominantly centered on the 16S rRNA gene, with limited or no support for comprehensive operon-level annotation. Greengenes, though historically popular, has not been updated in recent years, prompting the development of Greengenes2, which introduces phylogenetically consistent taxonomic annotations with improved genome integration. Greengenes2 unifies 21,074,442 sequences sourced from the Living Tree project, the Earth microbiome project, and 16S rRNA sequence variants including full-length, near complete, and short V4 amplicon sequence variants (ASVs) into single–integrated reference tree [5, 8]. Similarly, MIrROR and GROND have quality-checked full-length 16S-ITS-23S rRNA operon sequences, which aids researchers in analyzing microbial community analysis. These databases aim to build on earlier efforts by incorporating more curated genomes and taxonomic clarity [6, 7]. Additionally, GROND also provides information on intragenomic variations within constituent regions and *rrn* operons. While *rrn*DB (ribosomal RNA operon copy number database) offers mainly rRNA gene copy number data [16], it majorly lacks 23S copy number data and does not have 5S copy number information at all, and does not include sequence alignments, secondary structures, or information on intragenomic variation. The major features of these databases, including their positive and negative aspects, have been outlined in Table 1. It must be noted that some of these databases such as RDP, while still referenced, are no longer regularly maintained and have become outdated in both sequence content and taxonomy. Additionally, many of these databases also suffer from incomplete taxonomic coverage and partially annotated entries, limiting their utility for full-spectrum microbial genomics. Even the most actively curated databases like SILVA fall short in offering integrated insights across all three rRNA genes (16S, 23S, and 5S), intragenomic identity variation, and structural predictions, all of which are increasingly relevant for accurate microbial classification, phylogenetics, and functional inference.

**Table 1:** Comparative table highlighting key features of major public databases dedicated to rRNA genes [3, 8, 16, 36, 37]. The table summarizes and contrasts various attributes across these databases, including the number of genome or sequence entries covered; availability of predicted or curated secondary structures; support for multiple sequence alignments and their visualization; inclusion of pairwise or percentage identity matrices; taxonomy-wise distribution and browsing; integration of all three rRNA genes (16S, 23S, and 5S); and genome-wide mapping or visualization of rRNA gene locations. This comparative analysis provides insight into the scope, strengths, and limitations of each database, emphasizing the need for integrated and comprehensive resources in rRNA-based microbial genomics.

| Category | Greengenes<br>2 | rrndb | SILVA | Mirror | GROND | 16S-23S-5S<br>rRNA<br>database |
| --- | --- | --- | --- | --- | --- | --- |
| Total genome |  | 48700 |  | 43653 | 571381 | 4974 |
| Total Sequences | 1,074,442<br>(16S only) |  | 10 million<br>(LSU &<br>SSU) | 97781 |  | (16S, 23S<br>and 5S) |
| Secondary structure | × | × | × | × | × | ✓ |
| Percentage identity | × | × | × | × | × | ✓ |
| Sequence alignments | ✓ | × | ✓ | × | × | ✓ |
| Taxonomy-wise<br>classification and<br>distribution | ✓ | ✓ | ✓ | ✓ | ✓ | ✓ |
| Data for 16S, 23S and 5S<br>rRNA | × | × | ✓ | × | × | ✓ |
| Chromosome location<br>details | × | × | × | × | × | ✓ |
| CIRCOS representation<br>of genomes | × | × | × | × | × | ✓ |
| Mapped NCBI sequence<br>viewer | × | × | × | × | × | ✓ |
| 16/23/5S Distribution<br>Per phylum | × | × | × | × | × | ✓ |

Despite the proliferation of all the above-mentioned databases, significant gaps remain in how rRNA gene content and organization are cataloged across bacterial and archaeal genomes. Most current resources emphasize large-scale rRNA sequence curation and taxonomy assignment [17], but do not systematically document how many copies of these three operon genes cumulatively are present in each genome, where they are located, how identical the intra-genomic copies are to each other, on which sites each copy has variations, and what are their putative secondary structures. This lack of detailed operon-level annotation and GCN information represents a major bottleneck in microbial genomics, particularly for researchers aiming to understand rRNA gene evolution, operon structure, and their implications for metagenomics and quantitative microbiome profiling. Intragenomic heterogeneity, a well-reported phenomenon [13, 14] is rarely accounted for in most databases. Moreover, such intragenomic heterogeneity information for 23S and 5S has been rarely documented. These gene copy number variations and their intragenomic heterogeneity have led to over- or under-estimation of microbial diversity in amplicon-based studies, potentially confounding ecological or evolutionary interpretations [18, 19]. Furthermore, extracting rRNA gene content from NCBI genome files (e.g., GBFF, GFF3, or RefSeq annotations) is a non-trivial task, especially for experimental researchers lacking bioinformatics expertise. Often, users must parse large archive files (e.g., SILVA’s ARB format), which are difficult to search and visualize. Overall, there is a clear need for a more accessible and biologically rich resource that bridges this usability gap. To address these limitations, we introduce the 16S-23S-5S rRNA Database, a comprehensive and user-friendly platform that consolidates sequence, alignment, structural, and copy number data for all three rRNAs across bacterial and archaeal genomes. This publicly available resource not only captures the taxonomic and genomic context of rRNA operon genes but also offers tools to explore intragenomic heterogeneity, sequence homology, and secondary structure predictions, empowering researchers to perform more accurate and nuanced microbiome analyses.

### Methodology for data collection and processing

Complete bacterial and archaeal genomes were retrieved from the NCBI RefSeq database as of February 20, 2024, restricted to those assemblies labeled as “Complete/Chromosome” and designated as Reference Representative under the RefSeq category [20]. The final dataset contained 4,974 genomes distributed into 4690 bacteria and 284 archaea. Each genome was mapped to its corresponding lineage in the NCBI Taxonomy database, which provides a hierarchical classification from superkingdom to strain level, enabling taxon-wise aggregation and analysis [21].

To identify ribosomal RNA (rRNA) genes, we utilized the RFAM database, which hosts manually curated multiple sequence alignments and covariance models for RNA families [22]. RFAM database harbors families associated with bacterial rRNA genes - RF00177 for the small subunit ribosomal RNA (SSU_rRNA_bacteria), RF00001 for the 5S ribosomal RNA (5S_rRNA), and RF02541 for the large subunit ribosomal RNA (LSU_rRNA_bacteria). For the identification and quantification of the rRNA genes, all rRNA gene sequences were extracted from the *rna_from_genomic.fna file for each genome was subjected to the hmmscan (HMMER 3.4) module against RFAM libraries [23]. After parsing all files, each sequence was assigned to its corresponding classes, and those high-quality rRNA sequences along, with their annotations, were extracted and further analyzed. This workflow enabled high-confidence annotation of rRNA genes, reducing errors from potential misannotations in public files.

To visualize the distribution of all three types of rRNA genes across bacterial and archaeal genomes, we generated Circos plots using Circos v0.69-8 (Perl 5.034000) [24], where 16S, 23S and 5S rRNA genes were marked in red, green, and black respectively. GC skew values were calculated using an in-house Python script with a window size of 5,000 bp and a step size of 1,000 bp, and it was shown as an additional track. To provide interactive genome navigation, we embedded the NCBI Sequence Viewer (v3.50.0) [25], which allows users to examine genomic architecture and features through options such as zoom, pan, search, region selection, and track toggling, enhancing usability for users with varying bioinformatics expertise.

To assess intragenomic heterogeneity among multiple copies of rRNA genes within each genome, we subjected each copy of 16S, 23S, and 5S rRNA genes within a genome to BLASTn (v2.15.0+) against the database made of each copy [26] and pairwise percentage identity was calculated. The BLAST results were parsed using custom Python scripts to generate identity matrices for each 16S, 23S, and 5S rRNA class per genome. These matrices serve as a proxy to detect intragenomic heterogeneity, where lower pairwise identity values suggest the presence of divergent rRNA gene copies within a genome, potentially affecting taxonomic resolution and functional interpretation.

For each genome, the extracted copies of 16S, 23S, and 5S rRNAs were aligned using Clustal Omega (v1.2.4) from the European Bioinformatics Institute (EBI) [27]. The alignments were visualized using MView, which renders HTML-based outputs with color-coded sequence conservation [28], enabling a clear representation of conserved and variable regions across rRNA copies.

To predict secondary structures, rRNA sequences were processed using R2DT, a template-based rRNA drawing tool that utilizes covariance models for structural inference [29]. The nucleotide color scheme in the output structures indicates alignment with the reference model: black denotes nucleotide identity, green represents substitutions, magenta marks insertions, and blue shows nucleotides repositioned relative to the template. These structures provide additional insight into the conserved architecture of rRNA genes and may aid in the interpretation of evolutionary and functional constraints at the structural level.

### Database content

The 16S-23S-5S rRNA database has been designed to offer a comprehensive, intuitive, and interactive platform for exploring the ribosomal rRNA sequence, alignment, and structure data across bacterial and archaeal genomes. The navigation bar serves as the primary access point for all major features, including a query tab for organism-specific searches, a string-based search function, a statistics tab summarizing database-wide trends for each 16S/23S/5S at the phylum level, a user manual with detailed instructions, download option to extract all data via a single click, and resource section showcasing similar tools and databases (Figure 1A).

**Figure 1:**
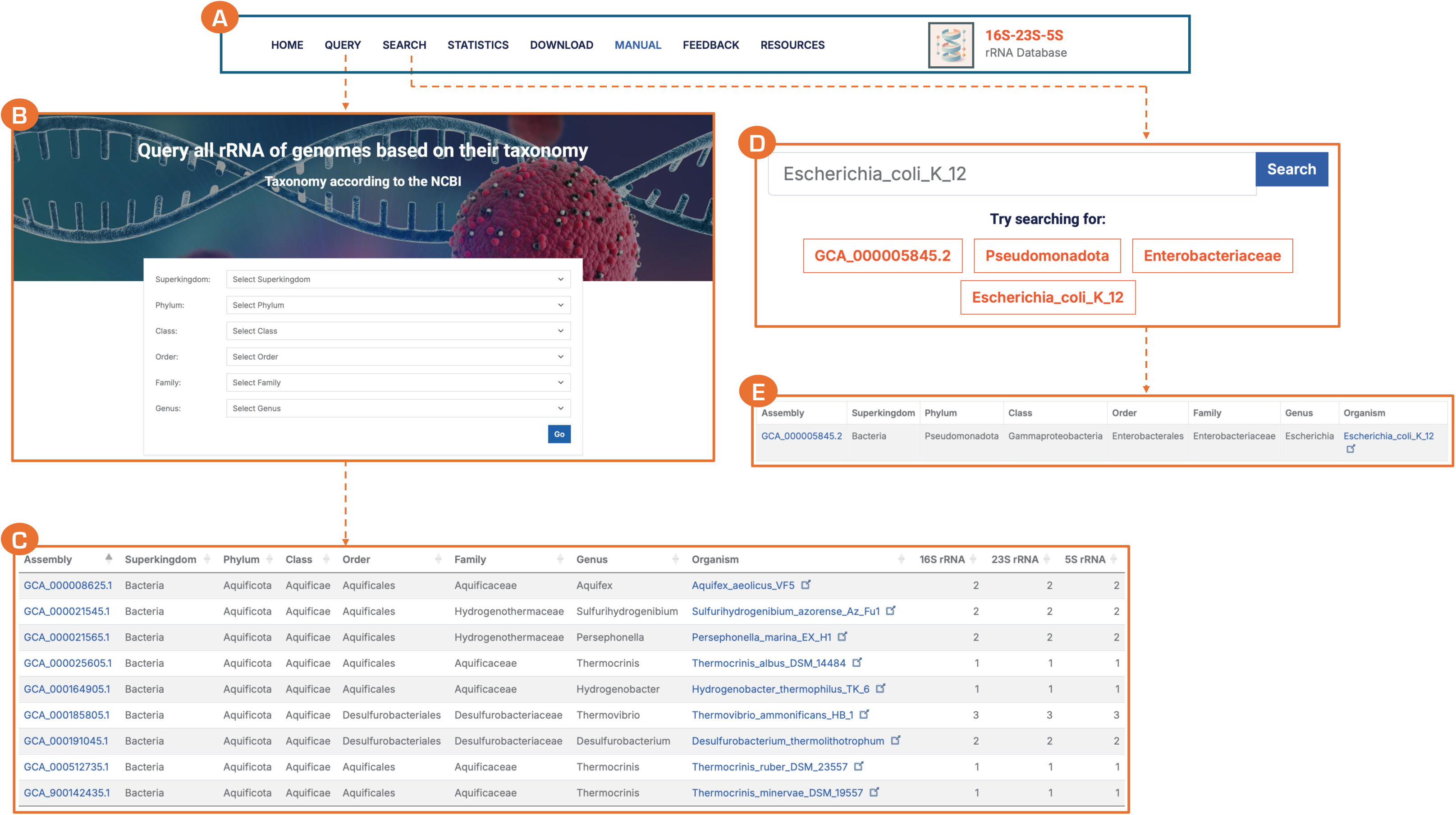
Search options in 16S-23S-5S rRNA database (A) Main navigation bar with query, search, statistics, other resources, and help tabs. (B) Users can query organism entry according to taxonomy using the pull-down menu (C) Word-based search option (D) Result for taxonomy-based search (E) Result for word-based organism entry

Users can access organism entries through a taxonomy-based drop-down menu, allowing them to build their query using different taxonomic ranks. Each higher taxa query provides a list of all lower-rank taxonomies. For example, the taxon page for Bacteria aggregates data across all bacterial classes, followed by all orders per class, all families per order, and all genera per family. For example, to access a narrower page for the genus *Myxococcus*, the user must build the full taxonomy query for this entry (Figure 1B). A sort-enabled table lists detailed metadata, including NCBI Assembly ID, taxonomic lineage (superkingdom to genus), organism name, and the copy numbers of 16S, 23S, and 5S rRNA genes (Figure 1C). In contrast to building a query, the search tab supports flexible keyword-based queries. Users can input specific assembly IDs such as “GCA_000005845.2” or taxonomic ranks such as “Pseudomonadota” or organism names (e.g., “Escherichia coli”) to access relevant entries (Figure 1D, 1E).

The detailed webpage of each entry has NCBI tax ID, NCBI assembly ID, taxonomy, genome size, number of chromosome and plasmids, GC%, number of genes, and number of tRNAs, as well as the 16S, 23S, and 5S rRNA counts (Figure 2A). It also includes hyperlinks to external NCBI Assembly ID and NCBI taxonomy web pages. It provides a valuable resource for investigating possible connections and associations among these genetic components.

**Figure 2:**
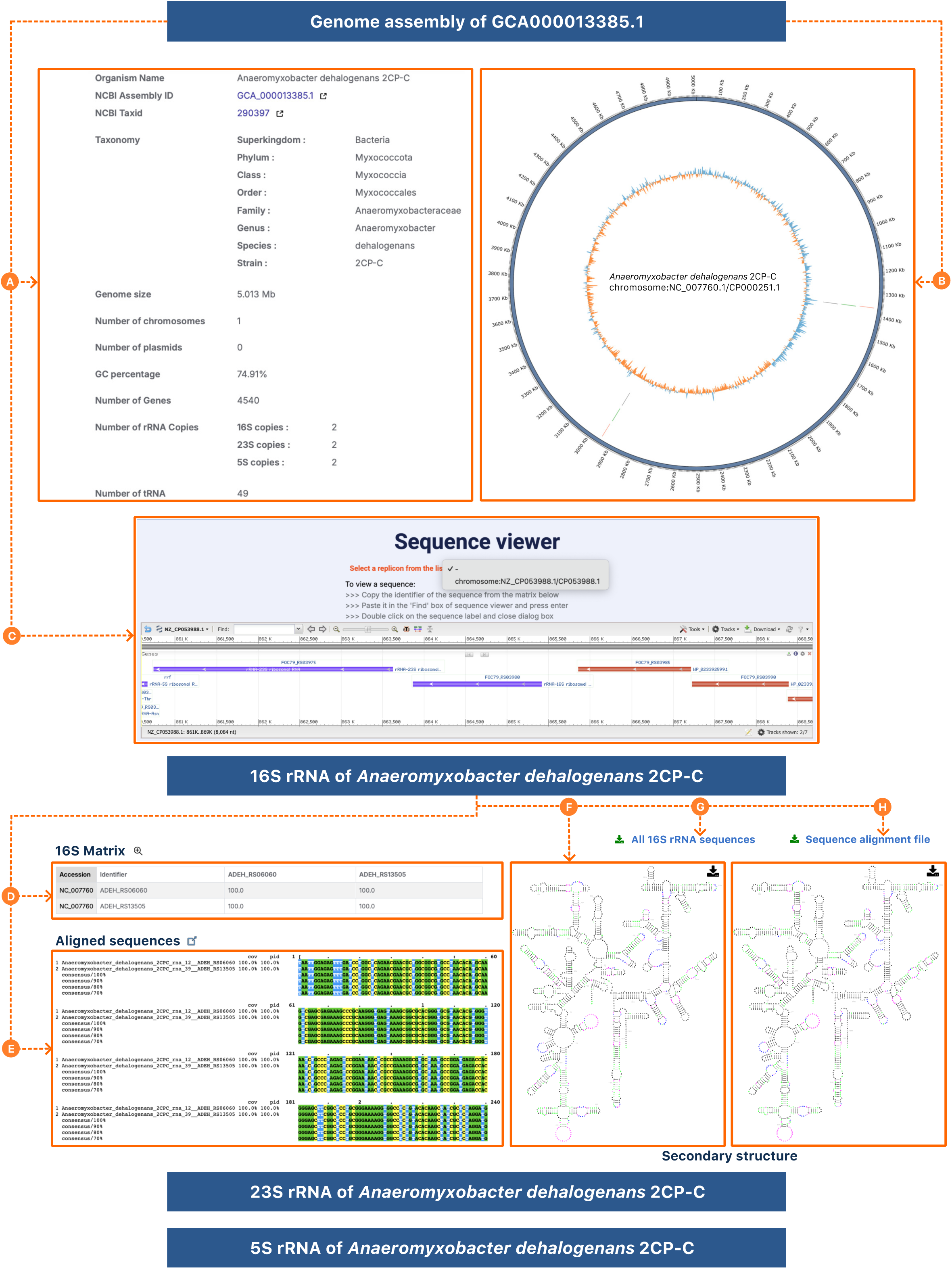
Database snapshots illustrating detailed organism page for *Anaeromyxobacter dehalogenans* 2CP-C (A) genome statistics panel displaying metadata such as the NCBI organism name, taxonomy ID with full taxonomic lineage, genome size, number of chromosomes and plasmids, GC content, total gene count, rRNA gene copy numbers, and tRNA count. (B) CIRCOS plot visualizes the complete genome structure, where the innermost ring represents GC skewness and rRNA genes are highlighted with distinct color codes (black for 5S, green for 23S, and red for 16S) as concentric bands progress outward. (D) embedded NCBI Sequence Viewer rendering the annotated 16S rRNA gene from *A. dehalogenans* 2CP-C, enabling interactive review of genomic coordinates. (D) predicted secondary structures for all 16S rRNA gene copies, aiding structural and functional interpretation (E) link to download all 16S sequences for organism (F) link to download sequence alignment file (G) percentage identity matrix depicting intragenomic similarity among 16S rRNA gene copies. (H) visualization of multiple sequence alignment of both 16S rRNA copies within the genome. All the above components collectively provide a granular exploration of rRNA gene diversity, organization, sequence heterogeneity, and structural attributes in any organism under study.

Circos plots of all genomes within an organism enable researchers to explore the 16S, 23S, and 5S rRNA gene localization across the chromosomes and other extrachromosomal replicons. As shown in Figure 2B, *Anaeromyxobacter dehalogenans* 2CP-C organism (NCBI Assembly ID GCA_000013385.1) harbors two copies of 16S, 23S, and 5S rRNA genes on its single chromosome (NC_007760.1) at around 1380 and 2950 Kb (<u>weblink</u>). In contrast, *Aliivibrio fischeri* ATCC 7744 (JCM 18803 or DSM 507) has two chromosomes and zero plasmids with 10 copies of 16S and 23S rRNA and 11 copies of 5S rRNAs. Circos plot depicts that 10 16S rRNA operons are present on the larger chromosome (NZ_CP092712), whereas one single copy is present on the smaller chromosome (NZ_CP092713) (<u>weblink</u>). Such plots also showcase multiple replicons and the relative presence and absence of rRNA subunits, such as in the case of *Bacillus cereus* FORC_047 (NCBI Assembly ID GCA_002220285.1), both plasmids do not have any rRNA subunit (<u>weblink</u>). However, *Azospirillum argentinense* Az39 (NCBI Assembly ID GCA_000632475.2) has one chromosome and five plasmids, which have two copies on the main chromosome, whereas seven copies are present on three out of five plasmids, which can be easily spotted via Circos plots (<u>weblink</u>).

The embedded NCBI sequence viewer further allows users to browse rRNA sequences in the sequence viewer, where they can visualize each gene, location, strand information, and their nearby gene architecture in a live manner. Users can select replicon accessions from the drop-down menu in the viewer to see specific replicon. To search for any identified 16S, 23S, and 5S rRNAs, users may search their locus tags from the percentage identity matrices (Figure 2C), present below the sequence viewer.

The subsequent section of this webpage is divided into three subsections, providing detailed information about each of the 16S, 23S, and 5S rRNAs **(Figure 2)**. To help users explore intragenomic heterogeneity per rRNA category within each genome, each organism entry has a percentage identity matrix for 16S, 23S, and 5S rRNA sequences with accession number and locus tags of respective genes **(Figure 2D)**. For example, *Caminibacter mediatlanticus* TB-2 (NCBI Assembly ID GCA_005843985.1), a member of the class Epsilonproteobacteria and an inhabitant of deep-sea hydrothermal vents (<u>weblink</u>), we observe four copies of each 16S, 23S and 5S rRNA sequences on its single chromosome. While three 16S rRNA copies are 100% identical to each other, one copy (FE773_RS00760) exhibits a substantial divergence, displaying only 82.99% identity to the others **(Figure 3A)**. Further analysis of its alignment reveals significant insertions within FE773_RS00760, potentially indicative of evolutionary events such as horizontal gene transfer or specialized adaptation to its extreme environment, as seen in other extremophiles with divergent 16S rRNA genes [30]. For 23S rRNA, too, three are identical, but one copy (FE773_RS00745) shows only 93.88% identity to others. However, all four copies of 5S rRNAs were 100% identical to each other, underscoring differential evolutionary pressures across the rRNA genes.

**Figure 3:**
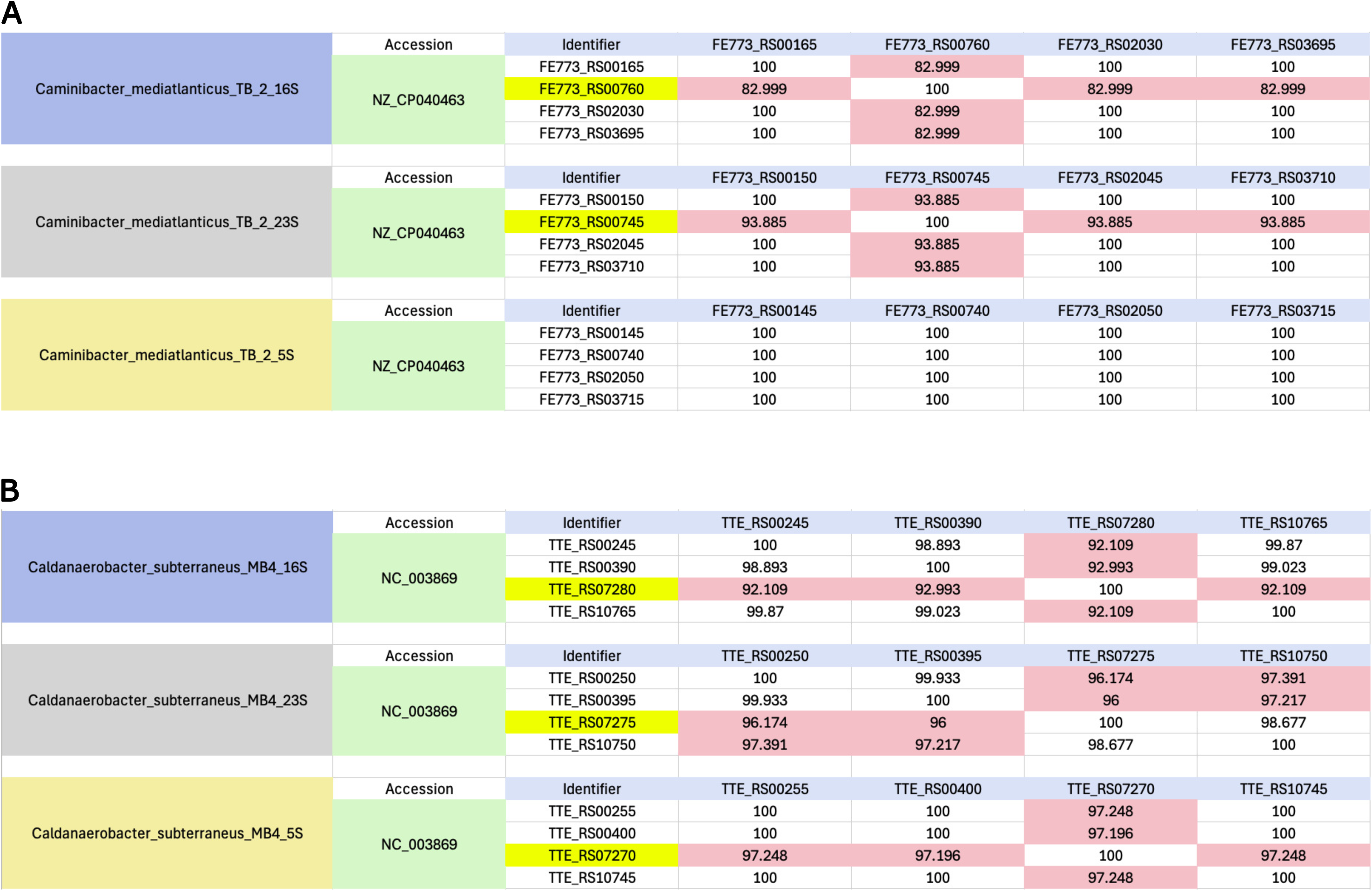
Cumulative percentage identity matrix illustrating intragenomic heterogeneity among 16S, 23S, and 5S rRNA gene copies within two representative genomes. Each cell represents the pairwise percentage identity between individual 16S, 23S, and 5S rRNA gene copies from the same genome, with pink shades indicating <98% sequence identity. The matrix highlights the range and distribution of intragenomic variation, emphasizing subtle but significant sequence divergence that may impact downstream phylogenetic and taxonomic analyses. The two examples shown correspond to the genomes (**A.** *Caminibacter mediatlanticus* TB-2 and **B.** *Caldanaerobacter subterraneus* MB4) as discussed in the main text.

Another illustrative example of intragenomic heterogeneity is *Caldanaerobacter subterraneus* subsp. tengcongensis MB4, a thermophilic anaerobe from the class Clostridia with NCBI Assembly ID GCA_000007085.1 (<u>weblink</u>), has four copies, out of which one 16S/23S/5S rRNA copy shows only 92, 96, and 97% identity respectively, whereas the other three show >98% identity with the other copies **(Figure 3B)**. These variations can arise from mechanisms like gene conversion, recombination, or evolution of mutational hotspots in specific variable regions [31]. Such pronounced intragenomic heterogeneity, particularly in the 16S rRNA gene, is a recognized phenomenon that can significantly impact phylogenetic inference and diversity estimations, often leading to overestimations of microbial diversity if not adequately accounted for [15]. Overall, such studies have the potential to decipher complex evolutionary dynamics underlying the rRNA operon, including gene duplication, concerted evolution, and adaptive divergence within the kingdom Bacteria and Archaea.

The colored version of the alignment can also be found on the page for each respective category of rRNA (Figure 2E). These alignments have been generated using MView 1.68 [28] and show the coverage and percentage identity values, too, with reference to one of the first rRNA sequences per the genomic location. It further showcases the consensus sequence at 70, 80, 90, and 100% identity levels, allowing the user to find the variable regions in an easier manner. Users can access and download all FASTA sequences of respective rRNAs from the given link (Figure 2G) and text files with aligned rRNA sequences through a link provided at the top of each section (Figure 2H).

Another important aspect that the 16S-23S-5S rRNA database covers is providing the predicted secondary structures for each 16S, 23S, and 5S rRNA sequences in each organism (Figure 2F). While nucleotide sequences are the foundation of phylogenetics and molecular evolution, rRNA molecules fold into complex, highly conserved secondary structures, which are crucial for ribosomal function and often evolve under strong selective pressure to maintain their intricate folding patterns. By visualizing predicted secondary structures, this database allows researchers to consider not just sequence similarity but also structural conservation, which can be highly valuable for resolving relationships among closely related species where sequence differences might be subtle, but subtle structural variations could be more informative. Identification and analysis of the covarying sites facilitated by structural information can provide stronger phylogenetic signals than sequence data alone, as it indicates a functional constraint on the molecule.

### Database statistics

The 16S-23S-5S rRNA database currently curates genomic information from 4,690 bacterial and 284 archaeal organisms, systematically organized within their respective taxonomies ranging from Kingdom to species and strains. This wide dataset spans over 51 distinct phyla, highlighting the broad phylogenetic diversity encapsulated for such a high-throughput database. Kingdom Bacteria has several highly represented phyla including Pseudomonadota (1,891 genomes), Actinomycetota (846 genomes), Bacillota (773 genomes), followed by Bacteroidota (392 genomes), Mycoplasmatota (125 genomes), and Campylobacterota (105 genomes). Within the phylum Archaea, phylum Euryarchaeota predominates with 205 genomes. The remaining 44 phyla contribute fewer than 100 species per phylum, accentuating the current sampling bias towards well-studied microbial groups, a recognized limitation in genomic databases.

Analysis of rRNA copy numbers for each of the 16S, 23S, and 5S rRNAs reveals significant variability across prokaryotic genomes. Within studies of 4,690 bacterial genomes, the number of 16S rRNA gene copies ranges from 1 to 37, 23S rRNA gene copies from 1 to 39, and 5S rRNA gene copies from 1 to 32 **(Figure 4)**. For the 284 archaeal genomes, copy numbers were generally lower, with 1 to 5 copies for both 16S and 23S rRNA genes, and 1 to 9 copies for the 5S rRNA gene.

**Figure 4:**
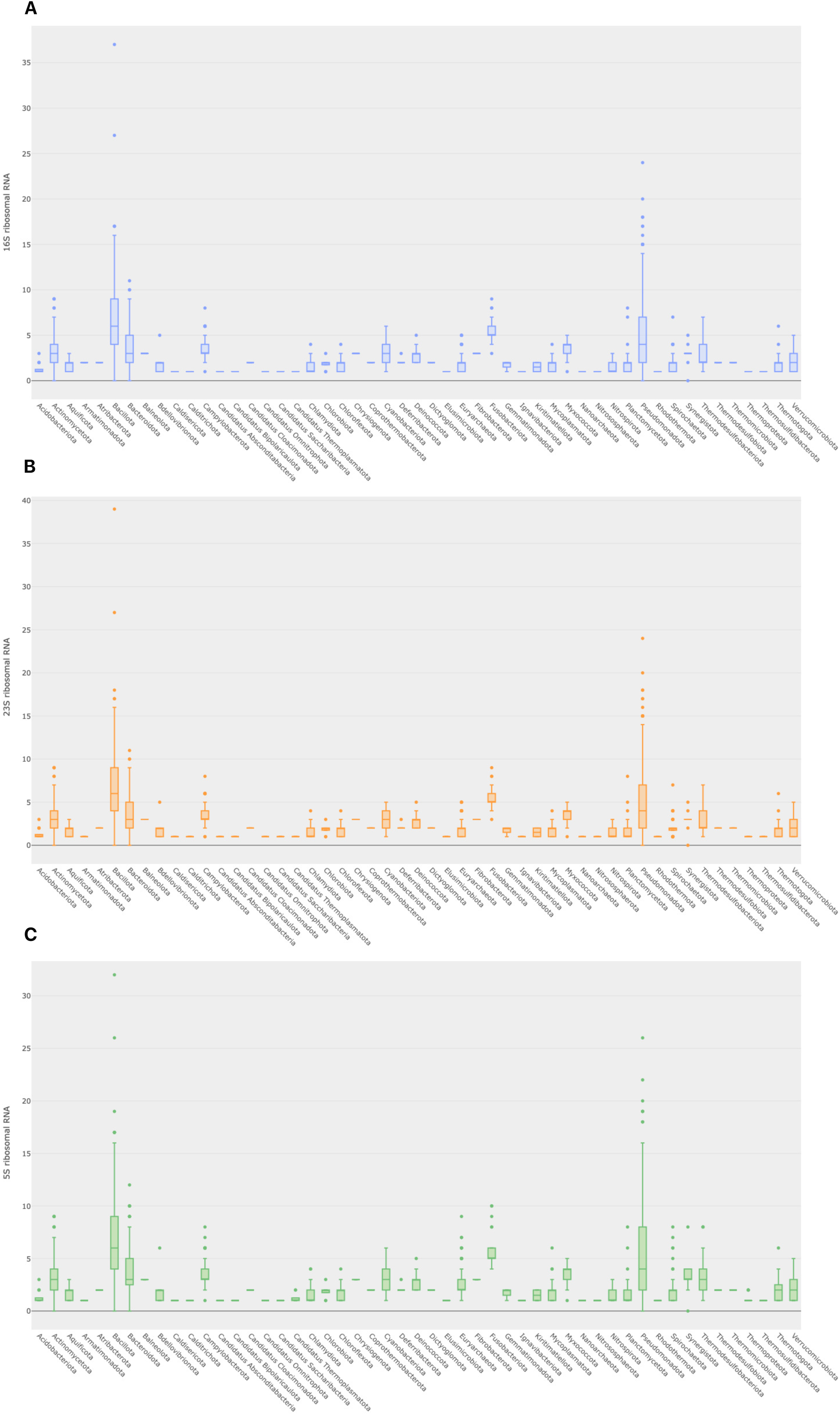
Phylum-wise distribution of rRNA gene copy numbers across bacterial and archaeal genomes, visualized using box plots. (A) Box plot showing the variation in 16S rRNA gene copy numbers across different phyla, indicating both median values and interquartile ranges. (B) Box plot representing the distribution of 23S rRNA gene copy numbers across phyla, generally mirroring the trends seen in 16S rRNA genes but with notable exceptions. (C) Box plot depicting the 5S rRNA gene copy number variation, which tends to be higher in number and more variable than the 16S and 23S counterparts in several phyla, which is also supported by their average copy numbers. These plots collectively highlight the taxonomic patterns in rRNA operon organization and the potential diversity of the ribosomal RNA gene.

The average rRNA gene copy numbers elucidate genomic strategies for efficient ribosomal synthesis. For bacterial genomes, the 16S rRNA gene exhibited an average copy number of 4.25±2.88, the 23S rRNA gene 4.24±2.89, and the 5S rRNA gene 4.44±3.08. This observed variability in rRNA gene copy numbers within bacteria has been previously correlated with the rate at which such bacteria respond to resource availability [32], with faster-growing copiotrophs, microorganisms that thrive best in nutrient-rich environments, typically possessing more copies [33]. In contrast, archaeal genomes displayed lower average copy numbers: 1.68±0.91 for both 16S and 23S rRNA genes, and 2.04±1.26 for the 5S rRNA gene.

Intriguingly, our dataset identifies 1,122 (22%) genomes where the number of 16S, 23S, and 5S rRNA copies are uneven. This phenomenon of uneven rRNA genes within operons, where gene ratios deviate from the canonical 1:1:1 stoichiometry, has been observed in several bacterial lineages and can arise from genomic rearrangements or incomplete operons, potentially impacting ribosomal biogenesis or reflecting specific evolutionary pressures [30, 34].

We also observed several phylum-specific rRNA gene copy numbers highlighting key differences. Such as within bacterial genomes, phylum Bacillota showcased the highest average rRNA gene copy numbers across all three rRNA genes (16S rRNA: 6.64±3.3; 23S rRNA: 6.63±3.3; 5S rRNA: 6.8±3.3). This high copy number in phylum Bacillota is consistent with their known fast-growth strategies allowing resource availability and adaptability to diverse niches, requiring rapid protein synthesis capabilities [33, 35]. Similarly, among the Archaea phyla, phylum Euryarchaeota members demonstrated the highest average rRNA gene copy numbers (16S rRNA: 1.94±0.9; 23S rRNA: 1.95±0.9; 5S rRNA: 2.4±1.3), reflecting the varying ribosomal demands within archaeal physiological diversity.

### Database construction

The 16S-23S-5S rRNA database was constructed with a focus on scalability, efficiency, and data integrity. The relational database schema, named “rrnadb”, was carefully designed and normalized to eliminate redundancy and optimize query performance. The relationships among the tables have been implemented via an entity-relationship diagram, which provides a structural overview of data organization. Some tables in the diagram, highlighted in cyan, are not currently implemented in the SQL database but are planned for future updates to accommodate scalability and increased data processing needs. PostgreSQL 14 was used to build this database, which was chosen owing to its robust support for relational data management, advanced indexing capabilities, and reliable transaction handling. While most data get stored within the database, the 16S, 23S, and 5S nucleotide sequences with their headers are presently retrieved from static .txt files instead of being directly stored in the database. This approach allows for a flexible transition, but as the database expands to incorporate larger datasets, the sequences will be stored and retrieved directly from SQL database tables.

The web interface was designed using a combination of HTML5, CSS3, JavaScript, and Bootstrap elements, ensuring an intuitive and responsive user experience. The backend web application was developed using Python’s Flask framework, providing a lightweight yet efficient server-side architecture. The application is hosted on a Ubuntu-based system and served using Apache HTTP Server (v2.4.54), ensuring reliable performance and seamless accessibility. Additionally, the database includes a mailer function integrated into the “Contact Us” page, which utilizes the SMTP protocol. This feature allows users to report bugs or provide suggestions directly to the authors, facilitating active feedback and continuous improvement. This structured approach ensures that the 16S-23S-5S rRNA database remains efficient, adaptable, and prepared for future enhancements, making it a valuable resource for researchers working with ribosomal RNA sequences.

### Future directions

We plan to include all chromosome and complete genome assemblies available in the NCBI RefSeq database in future updates. To improve user experience and ensure up-to-date data, we will regularly update the genome assemblies from NCBI RefSeq. In addition, the taxonomy from NCBI will be updated regularly. Furthermore, we plan to incorporate features related to rRNA operons, as well as their identification and annotation within genome assemblies. We also intend to provide new features that let users do on-site comparisons. These features and developments should enhance the platform’s utility for comparative genomic analysis.

## Conclusion

The 16S-23S-5S rRNA database represents a significant advancement in the study of prokaryotic ribosomal RNAs, moving beyond traditional single-gene approaches to offer a comprehensive user-friendly unified resource for rRNA sequence and structural analysis. By meticulously integrating sequence, predicted secondary structure, and crucial intragenomic heterogeneity data for over 4,900 bacterial and archaeal genomes, we address a long-standing void in microbial genomics infrastructure. The database currently houses approximately 62,000 rRNA sequences and further facilitates advanced research through its intuitive taxon and keyword-based search functionalities. Systematic integration of intragenomic heterogeneity, a phenomenon often overlooked but critical for accurate microbial classification, further distinguishes this resource, providing a robust framework to analyze variations that can influence microbial classification, shed light on the functional implications of rRNA polymorphisms, and contribute to a deeper understanding of prokaryotic genome evolution, particularly in relation to environmental adaptation and niche specialization. The inclusion of predicted secondary structures elevates the database beyond a mere sequence repository. This structural dimension enables researchers to explore the intricate interplay between sequence, structure, function, and evolution of these fundamental molecular machinery components. As the volume of genomic assemblies in NCBI RefSeq continues to expand, our database will be regularly updated to ensure its continued relevance and comprehensiveness. Future enhancements will focus on incorporating additional data and features, solidifying the 16S-23S-5S rRNA Database as an invaluable tool for taxonomists, microbial genomics, and microbiome researchers tackling challenges in the field of microbial genomics, systematics, metagenomics, and host-microbe interaction.

## Acknowledgments

We thank PARAM Seva, a National Supercomputing facility within IIT Hyderabad for their support in our research.

## Consent for publication

Not Applicable.

## Ethics approval and consent to participate

Not applicable

## Availability of data and materials

Authors have used open-source tools in this analysis, for which additional information has been provided at https://project.iith.ac.in/sharmaglab/rrnadatabase/.

## Competing interests

The authors declare no conflict of interest to disclose.

## Funding

GSha acknowledges the seed grant from IIT Hyderabad and the Start-up Research Grant (SRG) from the Science and Engineering Research Board (SERB) for supporting his research. JC and GShu thank the PhD fellowship support from the Department of Biotechnology (DBT, India) and the Center for Interdisciplinary Programs, IIT Hyderabad respectively.

## Authors’ contributions

GSha conceptualized the idea and supervised the project. JC performed the major analysis and wrote the first draft. GShu implemented the back- and front-end development for the web pages. JC, GShu, and GSha edited and finalized the manuscript. All authors approved the final version.

## Availability and Requirements

**Project name:** 16S-23S-5S rRNA database

**Project home page:** https://project.iith.ac.in/sharmaglab/rrnadatabase/

**Operating system(s):** Platform independent.

**Programming language:** Shell scripting, Javascript, HTML, CSS

**Other requirements:** Not Applicable

**License:** MIT

**Any restrictions to use by non-academics:** Not Applicable

## References

1. Woese CR. Bacterial Evolution. MICROBIOLOGICAL REVIEWS. 1987.

2. Van De Peer Y, Chapelle S, De Wachter R. A quantitative map of nucleotide substitution rates in bacterial rRNA. Nucleic Acids Research. 1996.

3. Quast C et al. The SILVA ribosomal RNA gene database project: Improved data processing and web-based tools. Nucleic Acids Res 2013;41. 10.1093/nar/gks1219

4. Cole JR et al. Ribosomal Database Project: Data and tools for high throughput rRNA analysis. Nucleic Acids Res 2014;42. 10.1093/nar/gkt1244

5. DeSantis TZ et al. Greengenes, a chimera-checked 16S rRNA gene database and workbench compatible with ARB. Appl Environ Microbiol 2006;72:5069–5072. 10.1128/AEM.03006-05

6. Seol D et al. Microbial Identification Using rRNA Operon Region: Database and Tool for Metataxonomics with Long-Read Sequence. Microbiol Spectr 2022;10. 10.1128/spectrum.02017-21

7. Walsh CJ et al. GROND: a quality-checked and publicly available database of full-length 16S-ITS-23S rRNA operon sequences. Microb Genom 2024;10. 10.1099/mgen.0.001255

8. McDonald D et al. Greengenes2 unifies microbial data in a single reference tree. Nat Biotechnol 2024;42:715–718. 10.1038/s41587-023-01845-1

9. Shajani Z, Sykes MT, Williamson JR. Assembly of bacterial ribosomes. Annu Rev Biochem 2011;80:501–526. 10.1146/annurev-biochem-062608-160432

10. Srivastava AK, Schlessinger D. MECHANISM AND REGULATION OF BACTERIAL RIBOSOMAL RNA PROCESSING. 1990.

11. Fleurier S et al. NAR Breakthrough Article rRNA operon multiplicity as a bacterial genome stability insurance policy. Nucleic Acids Res 2022;50:12601–12620. 10.1093/nar/gkac332

12. Johnson JS et al. Evaluation of 16S rRNA gene sequencing for species and strain-level microbiome analysis. Nat Commun 2019;10. 10.1038/s41467-019-13036-1

13. Sun DL et al. Intragenomic heterogeneity of 16S rRNA genes causes overestimation of prokaryotic diversity. Appl Environ Microbiol 2013;79:5962–5969. 10.1128/AEM.01282-13

14. Pan P et al. Microbial Diversity Biased Estimation Caused by Intragenomic Heterogeneity and Interspecific Conservation of 16S rRNA Genes. Appl Environ Microbiol 2023;89. 10.1128/aem.02108-22

15. Pei AY et al. Diversity of 16S rRNA genes within individual prokaryotic genomes. Appl Environ Microbiol 2010;76:3886–3897. 10.1128/AEM.02953-09

16. Stoddard SF et al. rrnDB: Improved tools for interpreting rRNA gene abundance in bacteria and archaea and a new foundation for future development. Nucleic Acids Res 2015;43:D593–D598. 10.1093/nar/gku1201

17. Almeida A et al. Benchmarking taxonomic assignments based on 16S rRNA gene profiling of the microbiota from commonly sampled environments. Gigascience 2018;7. 10.1093/gigascience/giy054

18. Louca S, Doebeli M, Parfrey LW. Correcting for 16S rRNA gene copy numbers in microbiome surveys remains an unsolved problem. Microbiome 2018;6. 10.1186/s40168-018-0420-9

19. Gao Y, Wu M. Accounting for 16S rRNA copy number prediction uncertainty and its implications in bacterial diversity analyses. ISME Communications 2023;3. 10.1038/s43705-023-00266-0

20. Goldfarb T et al. NCBI RefSeq: Reference sequence standards through 25 years of curation and annotation. Nucleic Acids Res 2025;53:D243–D257. 10.1093/nar/gkae1038

21. Schoch CL et al. NCBI Taxonomy: A comprehensive update on curation, resources and tools. Database. 2020. Oxford University Press, 2020, 2020

22. Ontiveros-Palacios N et al. Rfam 15: RNA families database in 2025. Nucleic Acids Res 2025;53:D258–D267. 10.1093/nar/gkae1023

23. Eddyl SR. A NEW GENERATION OF HOMOLOGY SEARCH TOOLS BASED ON PROBABILISTIC INFERENCE.

24. Krzywinski M et al. Circos: An information aesthetic for comparative genomics. Genome Res 2009;19:1639–1645. 10.1101/gr.092759.109

25. Rangwala SH et al. Accessing NCBI data using the NCBI sequence viewer and genome data viewer (GDV). Genome Res 2021;31:159–169. 10.1101/gr.266932.120

26. Camacho C, et al. BLAST+: Architecture and applications. BMC Bioinformatics 2009;10. 10.1186/1471-2105-10-421

27. Sievers F et al. Fast, scalable generation of high-quality protein multiple sequence alignments using Clustal Omega. Mol Syst Biol 2011;7. 10.1038/msb.2011.75

28. Brown NP, Leroy C, Sander C. MView: a web-compatible database search or multiple alignment viewer.

29. Sweeney BA et al. R2DT is a framework for predicting and visualising RNA secondary structure using templates. Nat Commun 2021;12. 10.1038/s41467-021-23555-5

30. Coenye T, Vandamme P. Intragenomic heterogeneity between multiple 16S ribosomal RNA operons in sequenced bacterial genomes. FEMS Microbiol Lett 2003;228:45–49. 10.1016/S0378-1097(03)00717-1

31. Liao D. Gene conversion drives within genic sequences: Concerted evolution of ribosomal RNA genes in bacteria and archaea. J Mol Evol 2000;51:305–317. 10.1007/s002390010093

32. Klappenbach JA, Dunbar JM, Schmidt TM. rRNA Operon Copy Number Reflects Ecological Strategies of Bacteria. APPLIED AND ENVIRONMENTAL MICROBIOLOGY. 2000.

33. Roller BRK, Stoddard SF, Schmidt TM. Exploiting rRNA operon copy number to investigate bacterial reproductive strategies. Nat Microbiol 2016;1. 10.1038/nmicrobiol.2016.160

34. Brewer TE et al. Unlinked rRNA genes are widespread among bacteria and archaea. ISME Journal 2020;14:597–608. 10.1038/s41396-019-0552-3

35. Lin Q et al. Microbial life strategy with high rRNA operon copy number facilitates the energy and nutrient flux in anaerobic digestion. Water Res 2022;226. 10.1016/j.watres.2022.119307

36. Szymanski M et al. 5SRNAdb: An information resource for 5S ribosomal RNAs. Nucleic Acids Res 2016;44:D180–D183. 10.1093/nar/gkv1081

37. Cabezas MP, Fonseca NA, Muñoz-Mérida A. MIMt: a curated 16S rRNA reference database with less redundancy and higher accuracy at species-level identification. Environ Microbiome 2024;19:88. 10.1186/s40793-024-00634-w

